# SARS-CoV-2 Virus like Particles produced by a single recombinant baculovirus generate potent neutralizing antibody that protects against variant challenge

**DOI:** 10.1101/2021.11.29.470349

**Authors:** Edward Sullivan, Po-Yu Sung, Weining Wu, Neil Berry, Sarah Kempster, Deborah Ferguson, Neil Almond, Ian M. Jones, Polly Roy

## Abstract

The Covid-19 pandemic caused by SARS-CoV-2 infection has highlighted the need for the rapid generation of efficient vaccines for emerging disease. Virus-like particles, VLPs, are an established vaccine technology that produces virus-like mimics, based on expression of the structural proteins of a target virus that can stimulate strong neutralizing antibody responses. SARS-CoV-2 is a coronavirus where the basis of VLP formation has been shown to be the co-expression of the spike, membrane and envelope structural proteins. Here we describe the generation of SARS-CoV-2 VLPs by the co-expression of the salient structural proteins in insect cells using the established baculovirus expression system. VLPs were heterologous ∼100nm diameter enveloped particles with a distinct fringe that reacted strongly with SARS-CoV-2 convalescent sera. In a Syrian hamster challenge model, a non-adjuvanted VLPs induced neutralizing antibodies to the VLP-associated Wuhan S protein, reduced virus shedding following a virulent challenge with SARS-CoV-2 (B.1.1.7 variant) and protected against disease associated weight loss. Immunized animals showed reduced lung pathology and lower challenge virus replication than the non-immunized controls. Our data, using an established and scalable technology, suggest SARS-CoV-2 VLPs offer an efficient vaccine that mitigates against virus load and prevents severe disease.

## INTRODUCTION

Severe Acute Respiratory Syndrome Coronavirus 2 (SARS-CoV-2) is the etiological agent of Coronavirus Disease 2019 (Covid-19). Since its initial identification in Wuhan, China, the virus has spread worldwide resulting in, to date, over 250 million confirmed cases and 5 million deaths. Genetic analysis of SARS-CoV-2 has shown that it shares 79.5% sequence homology with SARS, which emerged in 2003 (1).

SARS-CoV-2 is a member of the *Betacoronavirus* genus within the family *Coronaviridae*. The particle consists of a single copy of the 29.9kb positive sense single stranded RNA genome associated with the viral nucleocapsid protein (N) which is surrounded by an envelope derived from the host cell plasma membrane, gained at the time of virus budding, embedded with 3 further viral structural proteins, the Spike (S), Membrane (M) and Envelope (E) proteins (2). In addition to the structural proteins, coronaviruses express a large number of non-structural proteins, essential for replication but not present in the virus particle (3).

The most abundant of the Coronavirus (CoV) envelope-associated proteins is the M protein, a triple membrane-spanning glycoprotein that controls the conformation of the viral envelope (4). The M protein oligomerizes to form lattices on the membranes of the ER-Golgi intermediary compartments, resulting in membrane distortion (3). In addition, M directly interacts with the other coronavirus structural proteins enabling their recruitment into the nascent virus particle, such that the M protein is considered the primary driver of coronavirus assembly (2, 5). The envelope associated E protein is a small, 12kDa protein, which some studies have shown to be a viroporin (5). Very little E is incorporated into nascent virions, the majority remaining the ER and Golgi (6) but its absence impairs virus budding and leads to an attenuated phenotype (5, 7, 8). Co-expression of E and M in isolation is sufficient to cause the formation and release of virus-like particles (VLPs) (5, 9, 10), which can also incorporate the S protein if it is expressed in the same cells (9, 11, 12). The trimeric S protein is a type 1 transmembrane glycoprotein responsible for receptor binding and membrane fusion (13). S binds to angiotensin conversion enzyme II (ACE2) on the host cell surface via the outermost S1 domain (14, 15) leading to serine protease TMPRSS2 dependent cell entry driven by the S2 domain (16). S is also the immunodominant antigen of SARS-CoV-2 and antibodies that block the interaction with S1 with ACE2 are neutralizing (17). All currently approved vector-based or RNA vaccines for Covid-19 rely solely on S for their protective responses (18).

Previously, we described VLPs for SARS-CoV following expression of the requisite structural proteins in insect cells using a recombinant baculovirus (9). VLPs offer multivalent presentation of the S protein to stimulate humoral immunity as well as uptake by antigen presenting cells for the generation of T-cell responses (19, 20) and VLP based vaccines for other viruses, produced by the baculovirus insect cell system, are currently in use, proving evidence of scalability and acceptability (21, 22).

Here we report the generation of SARS-CoV-2 VLPs based on the co-expression of S, M and E proteins in insect cells. Purified VLPs presented as heterogeneous vesicular structures which were strongly recognized by convalescent human sera. Small mammal immunization trials generated neutralizing antibody that protected against subsequent variant live virus challenge. Our data demonstrate that SARS-CoV-2 VLPs produced by an established and scalable technology are a feasible vaccine for Covid-19.

## MATERIALS AND METHODS

### Construction of recombinant baculoviruses

A codon optimized sequence encoding the S protein of the original Wuhan SARS-CoV-2 isolate (QHD43416.1) was obtained from Integrated DNA Technologies as described (23) and cloned in place if the resident SARS S protein in vector pACVC3-SARS-S-E (9), leaving the E open reading frame unchanged. Similarly, an optimized SARS-CoV-2 M gene, was purchased from Eurofins Genomics and cloned into the baculovirus expression vector pAcYB2 (24). A single baculovirus expressing the 3 proteins was generated by co-transfection of pACVC3-SARS-E-Covid19-S and pAcYB2-Covid19-M with *Bsu36I* linearized AcMNPV DNA (25).

### Expression and Purification of VLPs

*Spodoptera frugiperda* (Sf9) cells were used for generation of recombinant baculoviruses and for subsequent virus amplification. For production of VLPs *Trichoplusia ni* (*Tnao38)* cells (26) were used. Virus was grown at an MOI of 0.01 and infected cells were incubated for up to 6 days before harvesting the supernatant. Virus infection for protein expression and VLP assembly employed an MOI of 5 and incubation for 4 hours, after which the cells were pelleted and re-suspended in fresh culture medium. Infected cells were then incubated for a further 72 hours. The resultant supernatant was harvested, clarified by centrifugation twice at 4500 × g, 4°C for 20 minutes, and the VLPs then pelleted by centrifugation through a 25% sucrose cushion, at 100,000 × g, 4°C for 100 minutes. VLPs were re-suspended in PBS and purified by centrifugation on a 20-60% sucrose gradient at 100,000 × g, 10°C for 18 hours. The gradients were fractionated and the presence of S protein assessed by dot blot with CR3022 (Absolute Antibody). The pixel density in each dot was assessed using Image J (27).

### SDS-PAGE and Western blot

Samples were resolved on Bolt™ 4-12% Bis-Tris Plus gradient gels (Invitrogen). Gels were either stained with Coomassie brilliant blue or subjected to Western blot using appropriate antibodies: rabbit polyclonal anti-SARS-CoV-2 S antibody (Abcam ab272504) or rabbit polyclonal anti-SARS-CoV/CoV2 M antibody (Novus Biologicals NB100-56569).

### Transmission Electron Microscopy (TEM)

Peak fractions from the sucrose gradient, identified by western blot with an anti-S antibody, were diluted 4-fold in PBS and concentrated to 10% of their original volume using spin filters with a cut-off of 1MDa. Carbon coated formvar grids were floated on droplets of the concentrated samples for 5 mins followed sequentially by 5 mins on a droplet of water and then on 1% uranyl acetate. The grids were blotted and allowed to dry before examination in a JEOL 2100 Plus microscope operating at 200kV.

### ELISA test

For the tests of VLP antigenicity, 96-well Nunc Maxisorb plates (Thermo Fisher Scientific) were coated with 0.5μg of purified VLPs per well diluted in 50μl of carbonate coating buffer (15mM Na_2_CO_3_, 36mM NaHCO_3_, pH9.6) and incubated overnight at 4°C. The plates were washed with PBS containing 0.05% Tween-20 (PBST) and blocked with Sea Block blocking buffer (Thermo Fisher Scientific) for 1 h at room temperature. 2-fold serial diluted human sera in blocking buffer were added to the plates and incubated for 1 h at room temperature. After three washes, the bound antibody was detected with goat anti-human Ig (γ chain specific)-HRP conjugated secondary antibody (Thermo Fisher Scientific) diluted 1:10,000 in blocking buffer. The assay reaction was developed with the 1-step Ultra TMB-ELISA substrate (Thermo Fisher Scientific). The reaction was left to develop for 5 min and then stopped with 2M H_2_SO_4_. The optical density was determined at 405nm using a plate reader. A surrogate marker of the neutralizing antibody response following immunisation, the cPass™ SARS-CoV-2 Neutralization Antibody Detection Kit (Genscript, L00847), was used as directed to detect antibodies that bind competitively to the receptor binding domain (RBD) of SARS-CoV-2.

### Immunogenicity studies

Five female Golden hamsters were administered 300µL (10μg) of non-adjuvanted VLP preparation under the skin on two occasions, 4 weeks apart. A control group of 5 hamsters did not receive any treatment. Two weeks after the second immunization hamsters were challenged intranasally with 50µL (1.5×10^5^ IU) of SARS-CoV-2 isolate B1.1.7, equally distributed between the two nostrils. The B.1.1.7 virus stock, obtained from Public Health England at Porton Down, was provided after propagation to passage 3 on the Vero/hSLAM cell line and deep sequenced to confirm absence of attenuating mutations at the furin cleavage site. Following virus challenge, all animals were weighed twice daily (AM and PM) and monitored for clinical signs. Clinical scores were recorded using a range of criteria such as behavior, appearance (ruffled fur, piloerection), breathing response and ear position. One hamster from each group was euthanized at 2 days post-challenge, two further animals from each group at 10 days post-challenge and the final two from each group at 14 days post-challenge. Oral swabs were taken from each animal at days 0, 1, 2, 3, 4, 7, 10 and 14 into Virus Transport Medium (VTM) (Hanks balanced salt solution with 2% heat-inactivated fetal calf serum, penicillin/streptomycin, 0.5µg/mL amphotericin B) for RT-qPCR analysis and dry swabs were taken for Point of Care (LFD) testing. Nasal swabs were taken at days 4, 7, 10 and 14 post-challenge. All procedures were carried out in UK laboratory in accordance with Home Office licensed procedures. Statistical analyses were performed using Graph Pad Prism v.9 and SigmaPlot v12.5.

### qRT-PCR

Total nucleic acid was extracted from a 200µL sample using the MagnaPure24 (Roche) External Lysis Pathogen 200 protocol with elution into 50 µL. qRT-PCR was performed in triplicate, with 5 µL per reaction, with primers and probe targeting the envelope protein as described by Corman *et al* (28). Viral shedding data was expressed in International Units per mL (IU/mL) calibrated against the WHO RNA standard for SARS-CoV-2 RNA (NIBSC: 20/146).

### Culture from swabs

20µL of the VTM sample was added to duplicate cultures of VeroTMPRSS2 cells either neat or diluted 1:2 in VTM. After 3 days the supernatant was removed, and the cells fixed for 30 minutes with 4% neutral buffered formalin followed by staining with 0.5% methyl violet. Where cell death was observed the presence of SARS-CoV-2 in cell culture supernatant was verified by qRT-PCR as described above. A sample was considered positive by culture if at least one of the four replicate wells showed destruction of the cell monolayer and was positive for SARS-CoV-2 by qRT-PCR.

### Point of care testing

The BioSensor SARS-CoV-2 Ag Kit (Oxford Biosystems) was used in accordance with manufacturer’s instructions to process swab samples from hamsters. Sample were added to lateral flow buffer and applied to the device and the output scored according to the WHO scoring system (https://extranet.who.int/pqweb/vitro-diagnostics/performance-evaluation). The LFDs were imaged and the relative pixel density in the test to control lanes was measured using Image J (27).

### Pathology

Intact lungs were collected at post-mortem, fixed for 72 hours at room temperature in 10% neutral buffered formalin (Sigma) and embedded in paraffin wax (VWR) in accordance with standard histological processes. Four micron thick sections were mounted on poly-L-lysine coated slides (VWR), de-waxed with xylene (Thermo Fisher Scientific) and re-hydrated via graded ethanol:water solutions (Thermo Fisher Scientific). Hematoxylin and eosin sections were evaluated for pathological changes associated with disease and assigned a score for each variable by two veterinary pathologists, independently. The samples were scored blinded.

### Immunohistochemistry (IHC)

Immunohistochemical staining was performed using the Leica Bond Polymer Refine staining system (Leica Microsystems DS9800). Onboard de-waxing was performed in accordance with the standard Leica Bond protocol and staining undertaken using IHC Protocol F with the following adaptations: additional non-specific block prior to primary antibody incubation (10% normal horse serum (Biorad), 1x Casein (Vector Labs) in PBS) and extended hematoxylin staining time (10 minutes). Antibodies were diluted to their optimal staining concentration in Bond primary antibody diluent (Leica, AR9352) as follows: SARS-CoV-2 Spike protein: 1:1000 (Leica, AR9961) 30minutes, (Rabbit PAb 40150-T62-COV2-SIB, Stratech Scientific/SinoBiologicals), SARS-CoV-2 Nucleoprotein: 1:2000 (Leica, AR9961) 30minutes, (Mouse Mab 40143-MM05-SIB, Stratech Scientific/SinoBiologicals). For both antibodies antigen unmasking was undertaken using Heat Induced Epitope Retrieval (HEIR) solution 1 (Leica AR9961) for 30 minutes at 100°C.

## RESULTS

### Baculovirus expression of SARS-CoV2 S, M and E proteins results in VLP formation in insect cells

A recombinant baculovirus driving the expression of SARS-CoV-2 S and M proteins from the polyhedrin promoter and the SARS-CoV E protein from the p10 promoter was constructed as described (Figure 1). Individual recombinant viruses expressing S, M or M+E were also constructed. The SARS-CoV and SARS-CoV-2 E proteins share 97% homology and are functionally exchangeable for the purposes of VLP production. Moreover, only a low level of E is incorporated into virus particles and it is not required for the neutralizing antibody response (3, 29). Infection of *Spodoptera frugiperda* (*Sf*9) cells with each recombinant virus followed by immunoblotting at 3 days post infection confirmed the expression of each of the CoV structural proteins separately and all proteins together in the case of the VLP competent construction. The recombinant viruses expressing S alone and the VLP expressed two bands identified with an S specific antiserum raised against a C-terminal peptide with molecular weights consistent with the complete S and the S2 cleavage product that were absent in cells infected with the M and M+E recombinant viruses. S protein maturation in insect cells is consistent with furin cleavage as previously shown (30). The M and M+E recombinant viruses expressed a band reactive with an M specific serum that was also produced by the recombinant baculovirus expressing VLPs (Figure 2). Thus, the structural proteins required for SARS-CoV-2 VLP formation were produced by a single recombinant baculovirus as was previously shown for SARS-CoV (9). Formal demonstration of E expression was not possible due to the lack of a specific serum.

**Figure 1.**
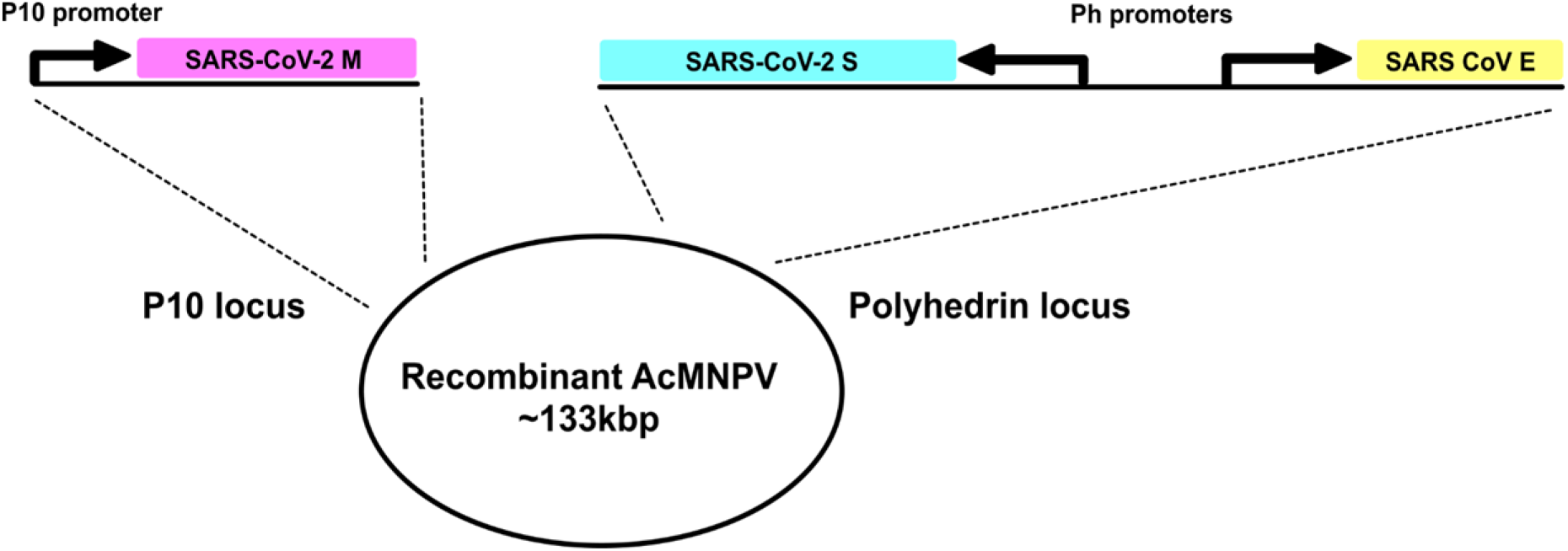
A schematic representation of the gene arrangements in the transfer vectors used to generate the recombinant baculovirus expressing SARS-CoV-2 VLPs. The SARS-CoV-2 M protein was expressed from the P10 promoter integrated at the P10 locus of the AcMNPV genome while S and E were expressed by back-to-back non-clashing polyhedrin promoters integrated at the polyhedrin locus of the genome.

**Figure 2.**
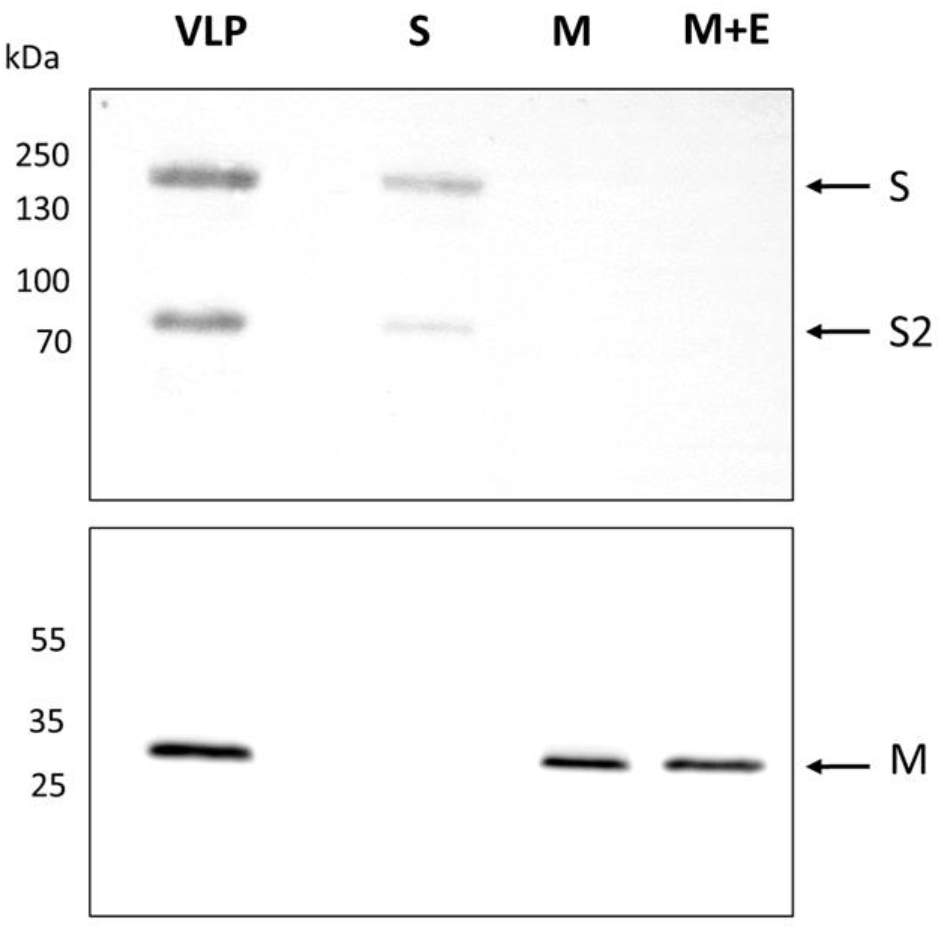
Expression of SARS-CoV-2 structural proteins by recombinant baculoviruses. Individual recombinant virus infections of *Sf*9 cells were harvested at 2 days post infection and total cell extracts analyzed by SDS-PAGE and Western blot with appropriate antibodies. The lanes are: VLP – virus expressing structural proteins S, M and E; S – virus expressing S only; M – virus expressing M only; M+E – virus expressing M and E. The upper panel was probed with an anti-S antibody and the lower panel probed with an anti-M antibody. Markers to the left of the gel are in kilodaltons.

To generate and purify VLPs, large scale infection (10^9^ cells) of the more productive *Tnao38* cells (26, 31) was performed and the infected cell supernatant harvested at 72 hours post infection. Cell debris was removed by low-speed centrifugation and VLPs remaining in the supernatant collected by ultracentrifugation. VLPs were further purified on a 20-60% sucrose gradient where they formed a distinct band at ∼35% sucrose (Figure 3A), the peak fraction of which reacted most strongly with anti-S monoclonal antibody CR3022 (Figure 3B) and showed 3 prominent bands at ∼180, 30, and 10 kDa, corresponding to S, M, and E proteins respectively on Coomassie blue stained SDS-PAGE (Figure 3C). Other proteins present at lower levels in the VLP preparation were insect cell membrane proteins present at the sites of VLP budding, as previously described by others (32).

**Figure 3.**
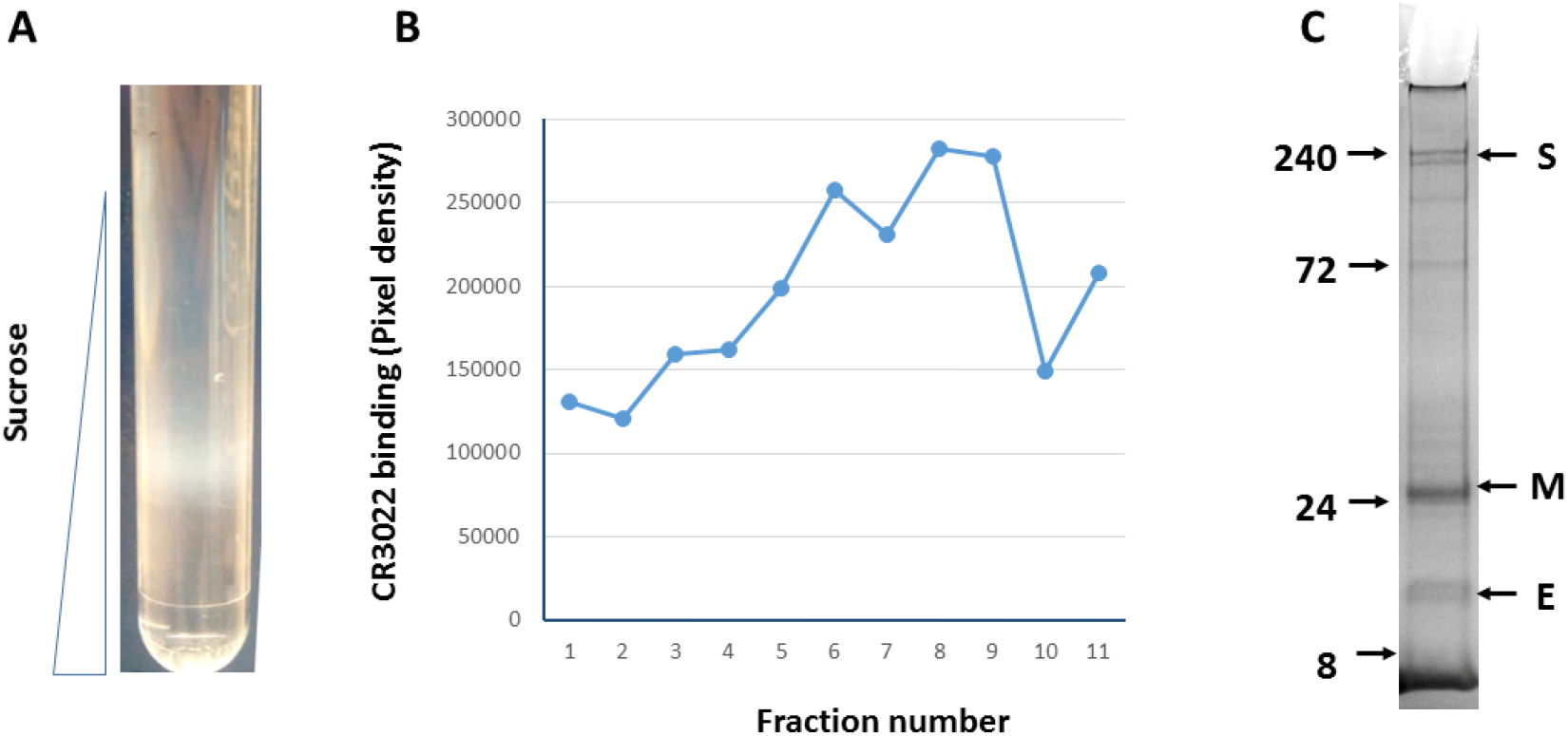
Purification of SARS-CoV-2 VLPs from infected *Tnao*38 cells. Infected cultures were processed as described and the resulting gradient (A) fractionated from the top. Each fraction was bot blotted to nitrocellulose, incubated with human anti-S monoclonal antibody CR3022 and developed with an anti-human HRP conjugate. The blot was scanned and the dot intensity recorded and plotted against fraction number (B). The peak fraction judged by appearance and immunoreactivity was analyzed by SDS-PAGE and stained with Coomassie brilliant blue R250 (C).

Fractions from the S monoclonal antibody strongly reactive ∼35% sucrose fraction were coated to TEM grids and analyzed by negative stain electron microscopy. Among the many heterogeneous vesicles present were vesicles of ∼100nm diameter that displayed a distinct fringe of ∼10nm in depth that were typical of the crown-like spikes which define coronavirus morphology (12, 33). Baculoviruses were absent from most fields of view consistent with their sedimentation to the bottom of the gradients used (Figure 4).

**Figure 4.**
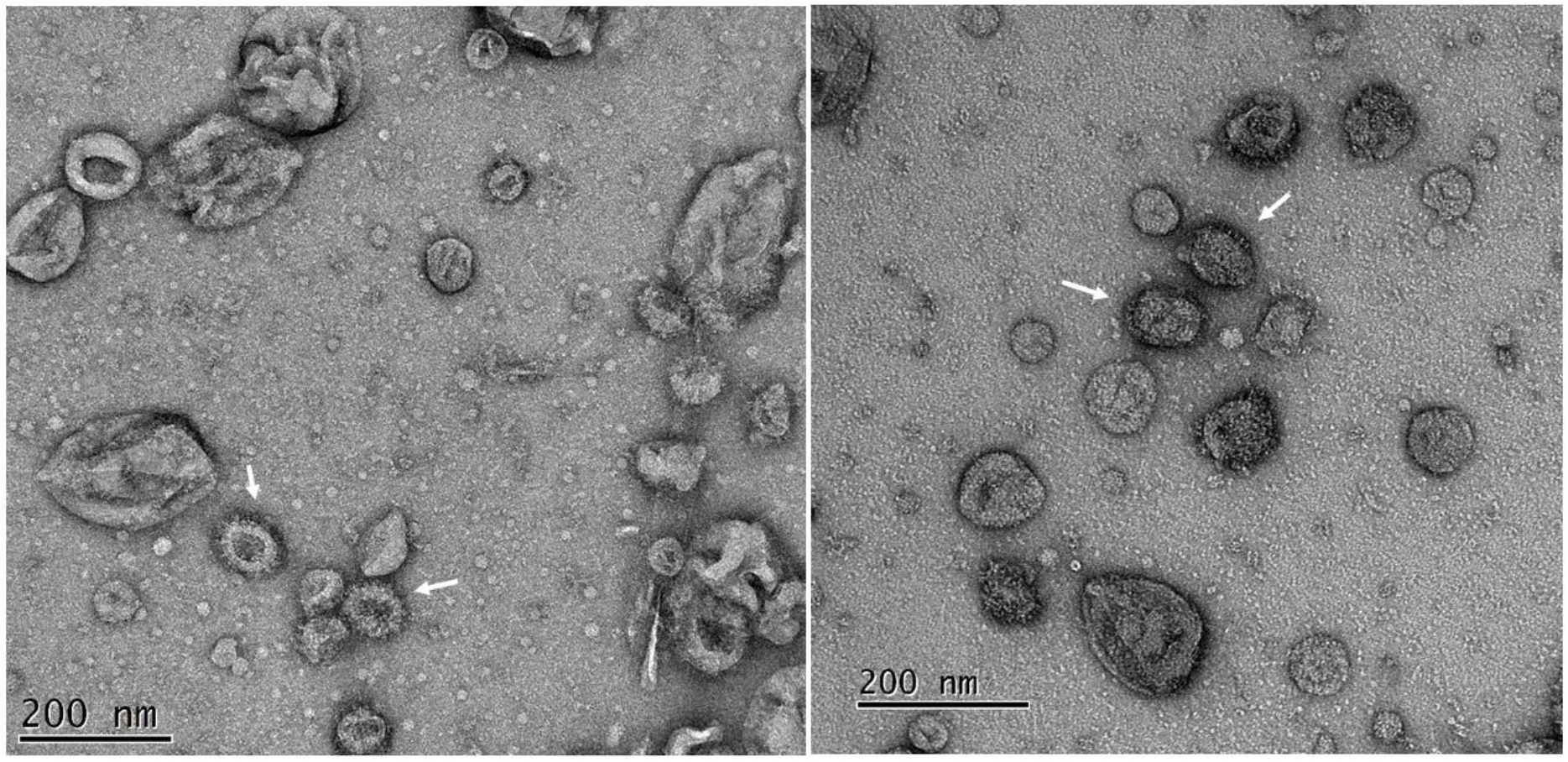
Transmission electron microscopy of purified SARS-CoV-2 VLPs purified from insect cells. Fractions were diluted to reduce the sucrose and re-concentrated before adsorption to carbon coated grids. The grids were strained with 2% uranyl acetate. Peak fractions (*cf*. Figure 3) of the VLPs showed multiple vesicle like structures among which are many with fringe like projections on their surface (arrowed). Two typical fields of the same adsorbed sample are shown.

### Antigenicity of SARS-CoV-2 VLPs

To assess relevant antigenicity, VLPs present in the peak fraction were coated to ELISA plates and probed with antibody positive convalescent sera previously screened using baculovirus expressed SARS-CoV-2 S1 protein (23). Multiple convalescent sera but not a naïve serum controls reacted strongly with immobilized VLPs (Figure 5A). For SARS-CoV-2 spike, extensive pepscanning of convalescent sera has shown that reactivity with linear epitopes is relatively poor (34, 35) suggesting that that VLPs used here display S in an antigenic form suitable for conformational antibody binding. To ensure that the sera used were not biased for insect cell expressed S1 protein and bound VLP displayed S preferentially, for example as a result of the high mannose glycan content of insect cell expressed proteins (36), VLP coated plates were also probed with a wholly different patient sera cohort pre-screened by peptide array (37). As before, reactivity was high for sera pre-scored as positive for infection but not for the serum pre-scored as negative (Figure 5B). Thus, insect cell derived SARS-CoV-2 VLPs are strongly antigenic.

**Figure 5.**
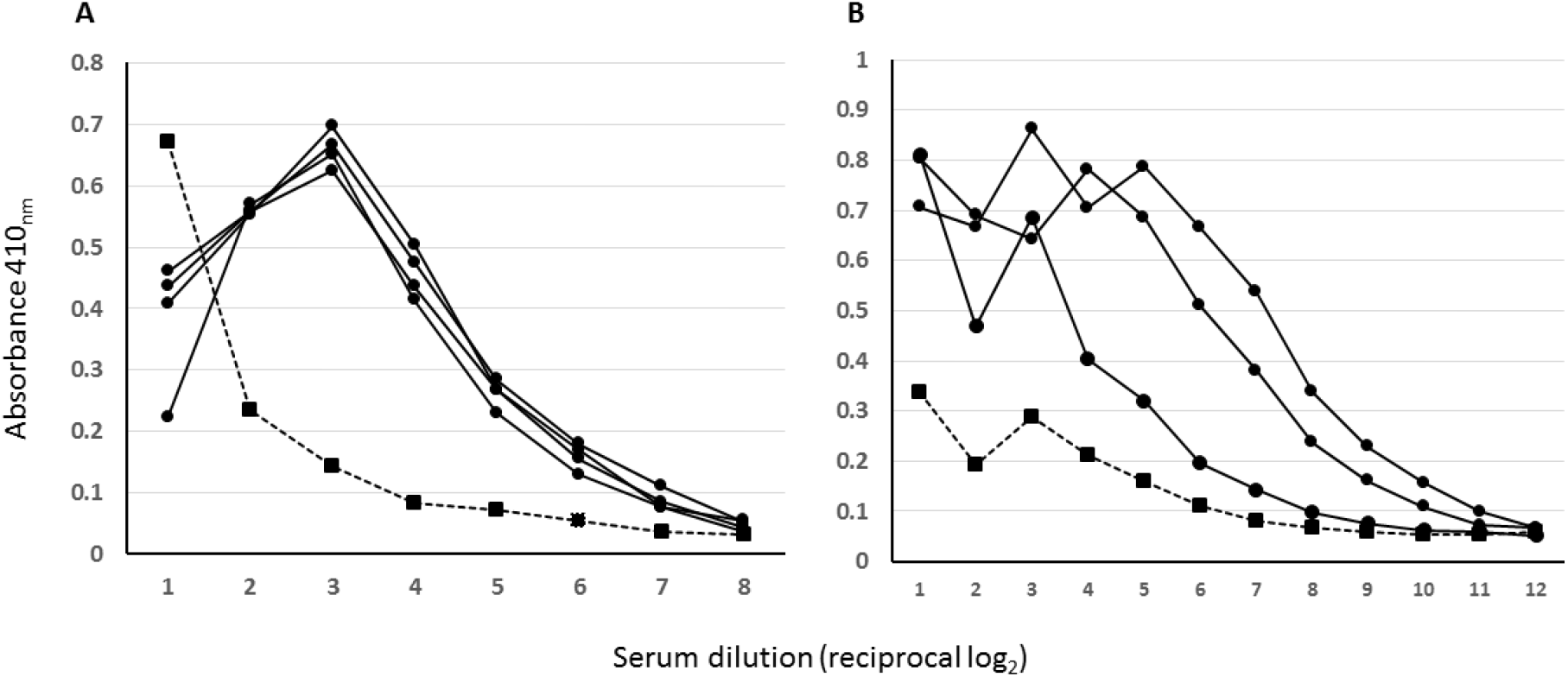
Antigenicity of SARS-CoV-2 VLPs determined by ELISA. Purified VLPs were adsorbed directly to the immunoblots overnight and blocked extensively before being probed with a twofold dilution series of pre-screened convalescent sera. Primary antibody binding was detected with an anti-human Ig HRP conjugate. A – Sera pre-screened by ELISA on purified SARS-CoV-2 S1 protein. B – Sera pre-screened by peptide array. In both assays filled circles and solid lines are sera that pre-screened positive while the single filled square dashed line is a negative

### Immunogenicity of VLPs in model animals

As the composition, appearance and immune recognition of the VLP preparation derived from insect cells infected with the VLP recombinant baculovirus was consistent with previous studies of CoV VLPs (12, 38, 39), insect cell derived VLPs bearing the SARS-CoV-2 S protein were assessed as a candidate vaccine in a hamster challenge model (40). Infection of Syrian hamsters with SARS-CoV-2 shows replication of the virus in the lungs with pathology similar to that reported for COVID-19 and a neutralizing antibody response that protects from subsequent infection (41, 42). Hamsters were vaccinated with gradient purified VLPs (n = 5, dose = 10μg per immunization) and boosted with the same material at 4 weeks post prime. Five additional control animals received no treatment. Seroconversion and the development of neutralizing antibody were tested using an anti-RBD neutralizing antibody surrogate competition immunoassay and was apparent in 4 out of 5 animals after the prime and in all animals in the immunized group after the boost (Figure 6A). No neutralizing activity was found in the control group.

**Figure 6.**
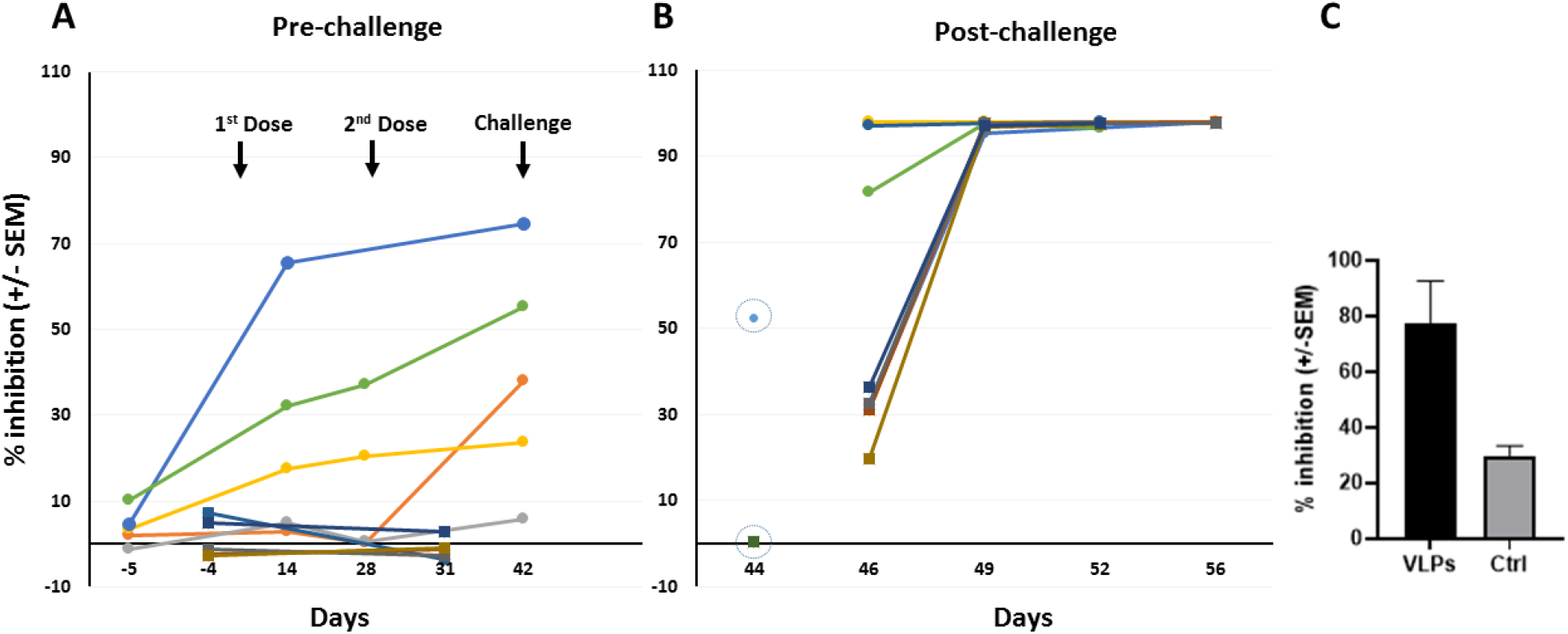
Neutralizing antibody development in immunized hamsters. Groups of Syrian hamsters were immunized with two doses of VLP candidate vaccine or untreated as controls and their sera tested for S binding using an RBD competition ELISA as described. A – Prior to live virus challenge. B - Post virus challenge. The dates of immunization and challenge are shown. Two animals, one from each group, culled two days after challenge are indicated at day 44 in the right panel. C – Serum titers to RBD determined in the two animals identified in B at the time of sacrifice.

Two weeks after the second immunization hamsters received a non-homologous intranasal challenge with a pre-titrated infectious dose of the SARS-CoV-2 B.1.1.7 variant and the infection monitored by virus recovery and virus antigen point of care lateral flow device. In addition, animals were weighed and scored for clinical signs as described (40). Neutralizing antibody titers rose within 4 days of challenge in the VLP immunized group and later also in the control group (Figure 6B). A single animal culled from each group at two days post challenge showed higher anti-RBD antibody titers in the VLP-immunized animal when compared with the control (Figure 6C).

The oral swabs of all animals from day 1 post challenge were positive for virus by genomic RNA RT-PCR but at day 7 post challenge were lower in the VLP immunized groups when compared with the controls (Figure 7A) and this difference was significant in the sampled nasopharyngeal swabs (p=0.044, one-tailed t-test) (Figure 7B). Virus antigen positivity by LFD of the oral swabs was strongly positive for both immunized and control groups at day 2, reduced at day 4, with a trend towards lower antigen levels in the immunized animals (Figure 7C (two animals from each group at day 2) & D (four animals from each group at day 4)) and absent by day 7 to the end of the experiment. Culturable virus was recovered from oral swabs throughout days 1 to 4 post-challenge with no discernible difference between the groups but not from any animal thereafter (not shown).

**Figure 7.**
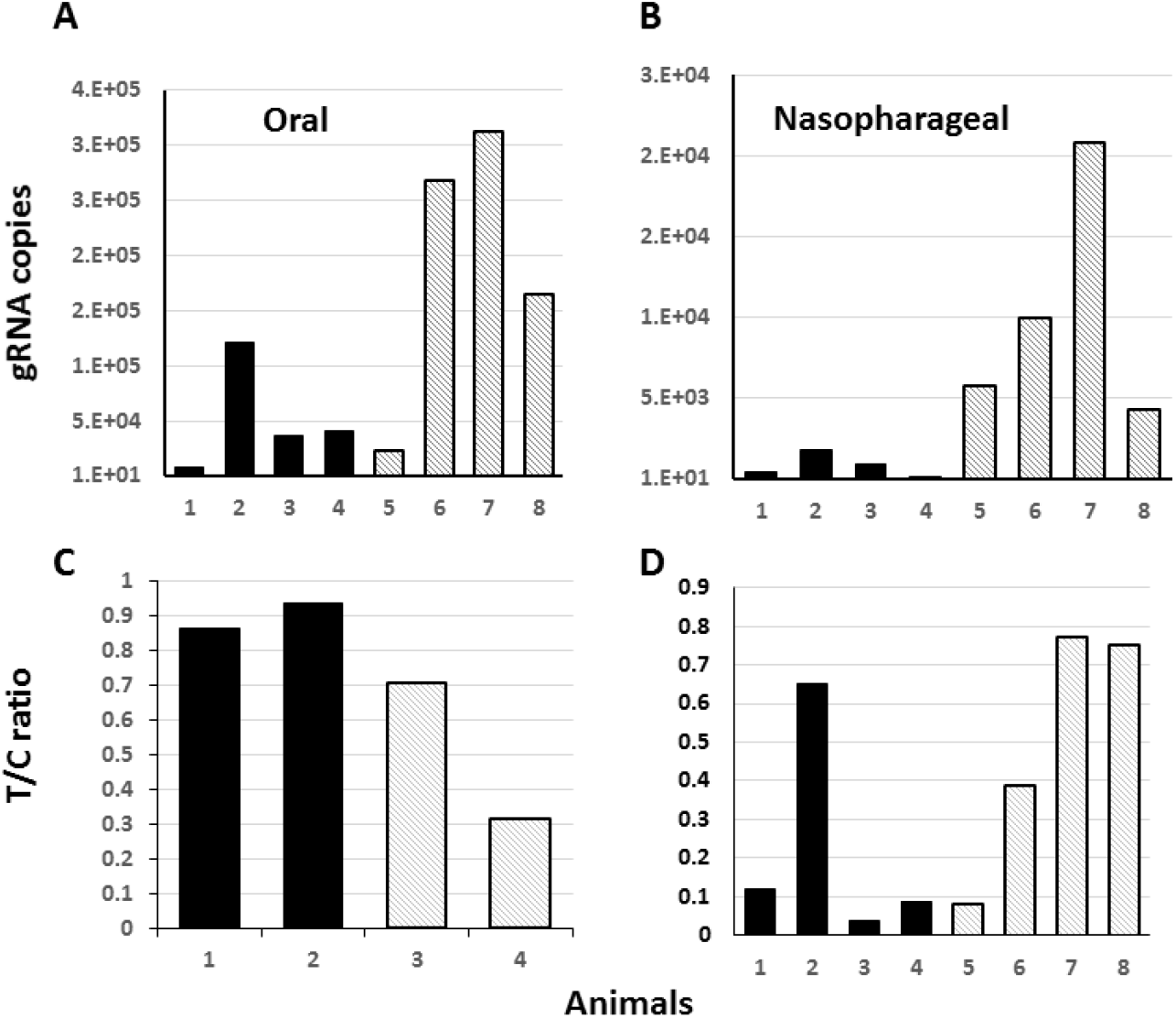
Presence of virus in animals following live virus challenge. Oral (A and C) or nasopharyngeal (B and D) washes were subjected to RT-PCR for genomic SARS-CoV-2 RNA (A and B) or point of care lateral flow device detection of virus antigen (C and D) and the loads recorded. The data shown is for day two (A and C) and day four post challenge (C and D). Only two animals from each group were tested by LFA at day 2. Neither assay was positive by day 7. Solid filled bars - vaccinated group. Stipple bars - control group.

Both groups lost weight for 5 days post challenge, but this was arrested in the VLP group from day 6 whereas the control group continued to lose weight for a further 3 days. Although all animals recovered by 14 days post challenge, the difference in weight loss between the groups at 5-8 days post challenge was statistically significant (p<0.05, ANOVA post-hoc Bonferroni) with the immunized animals recovering quicker (Figure 8).

**Figure 8.**
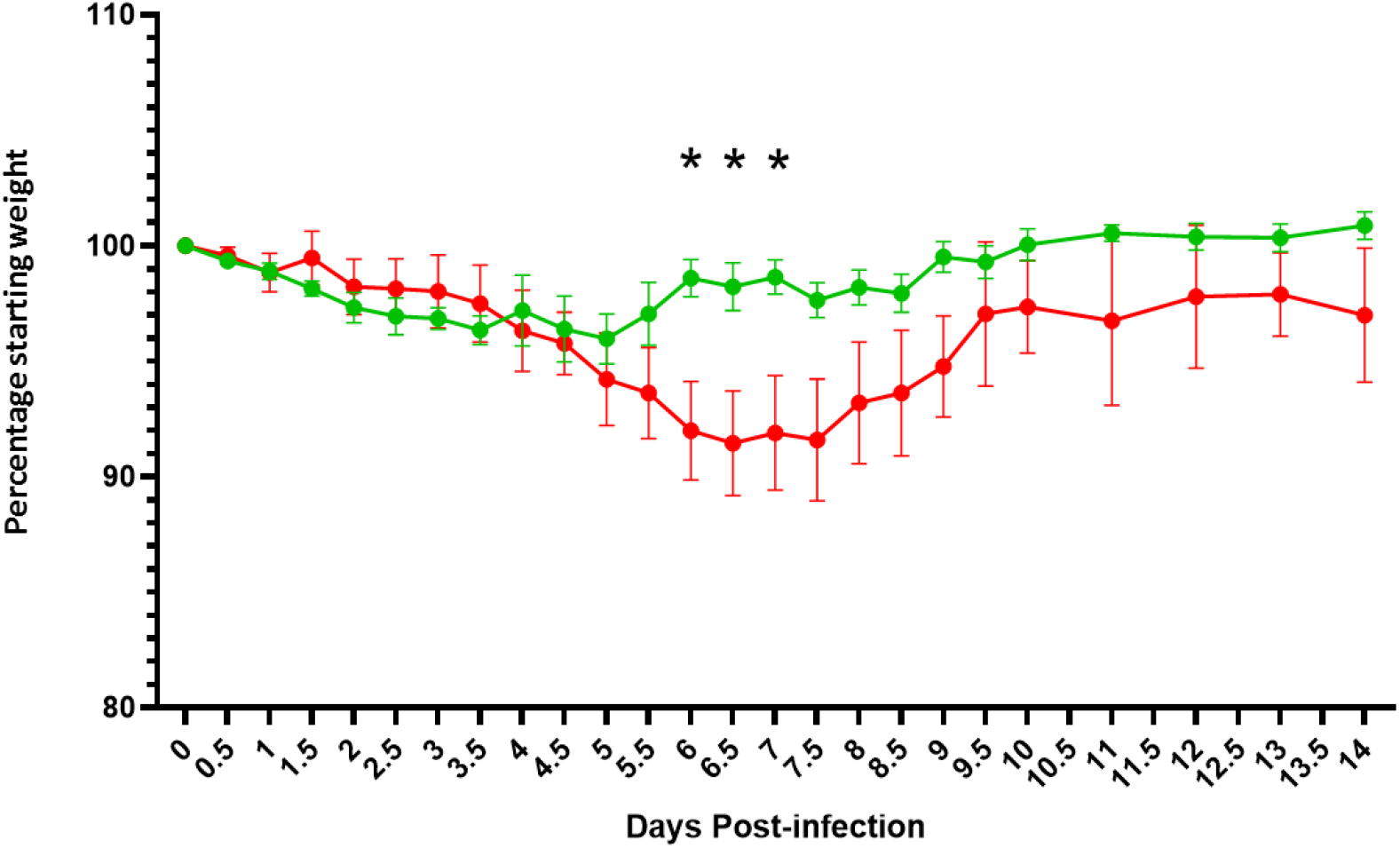
Average weight loss in both Syrian hamster groups, one immunized with VLP and the other control, following live virus challenge. Animals were weighed twice a day (AM and PM) and the average weight and variation among the group plotted. Significant deviation between the groups (p<0.05, ANOVA post-hoc Bonferroni) is indicated.

Pathology scoring of lung inflammation and disease revealed that the group that received the VLPs had markedly reduced bronchiolar and alveolar inflammation compared with control animals at 2 days post challenge, a feature which was retained to the end of the monitoring period at 10 days post-challenge (Figure 9 upper panels). Immunohistochemistry for the presence of the S protein in inflamed lung tissue showed clumps of S positive syncytial cells were present in both groups but that the overall level of S protein was much higher in the non-vaccinated animals when compared to those immunized with the VLPs at both of the time points tested (Figure 9 lower panels).

**Figure 9.**
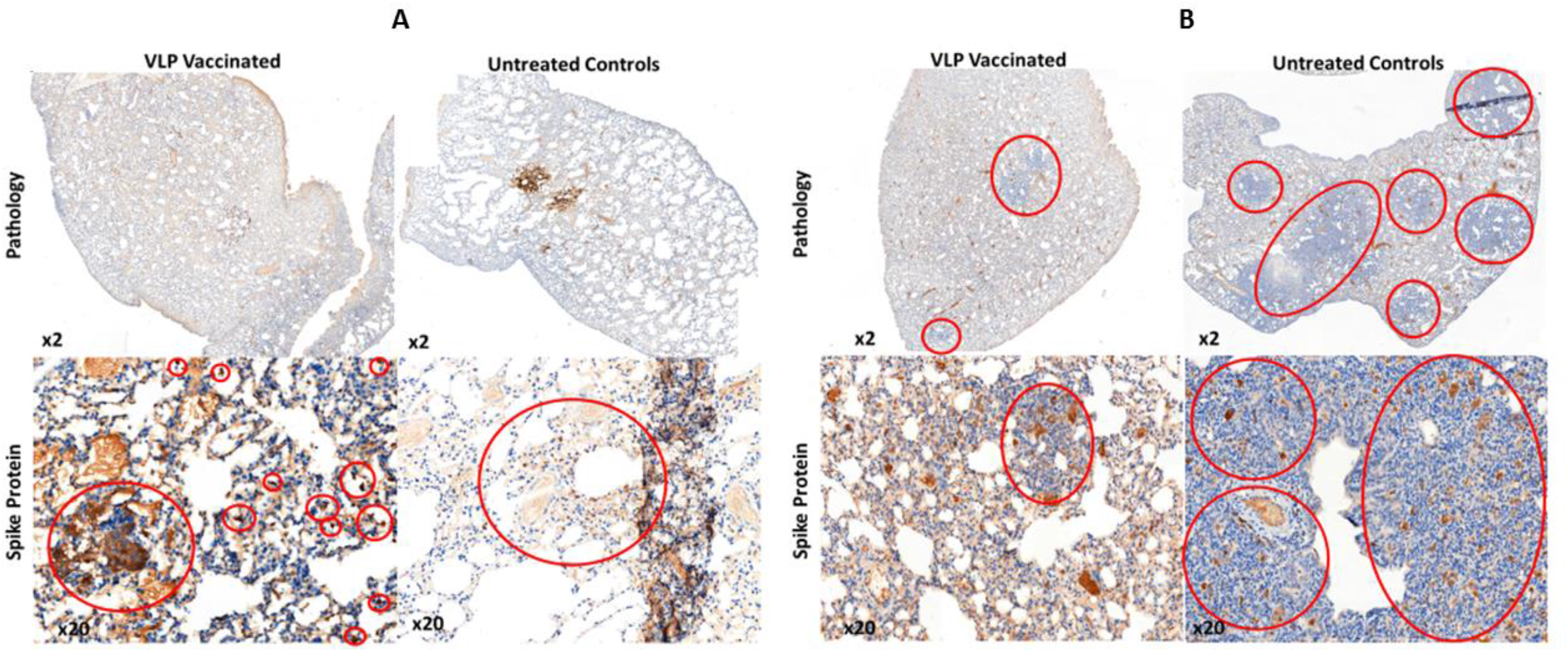
Histology of lung tissue taken from culled animal at 2 days (A) and 10 days (B) post live virus challenge. Upper panels - gross pathology revealed by H+E staining showing levels of lung sections showing vacuolization and eosinophilia (blue). Lower panels – immunohistochemistry with an anti-S antibody (brown).

## Discussion

The emergence of SARS-CoV-2 into the human population, most likely by spill over from a zoonotic reservoir (43), has led to the rapid development and licensing of several efficacious vaccines including those based on inactivated virus, adenovirus vectors, or direct nucleic acid (18, 44, 45). Other experimental vaccines, including VLP based vaccines have also been reported (38, 46, 47). Based on a previously described VLP of SARS CoV (9) we report here the development and test of a VLP for SARS-CoV-2 based on the co-expression of the S, M and E proteins using the baculovirus expression system. VLPs were shown to consist of the requisite CoV structural proteins, which were assembled into vesicle-like structures as shown by EM analysis. Further, these VLPs reacted with SARS-CoV-2 infected patient sera from two different serum sets, confirming their antigenicity and the combination of Western blot positivity, visualization and antigenic reactivity led to their use in trial immunizations. As an immunogen, non-adjuvanted SARS-CoV-2 VLPs generated a serum response in all immunized animals that competed with a standard RBD antibody-S protein interaction, consistent with neutralizing activity, which was boosted further following live virus challenge. No adverse events were recorded following immunisation consistent with extensive previous tests of insect cells expressed material as vaccines (22, 48). Immunisation with the SARS-CoV-2 VLPs did not prevent replication of the challenge virus but virus titers in the oral cavity, and particularly in the nasopharyngeal samples, were reduced when compared with the control group. The SARS-CoV-2 VLPs used here were assembled using the Wuhan S protein but the challenge SARS-CoV-2 virus stock was the B1.1.7 S variant that carries the N501Y residue change, which has been reported to improve ACE-2 binding and partly evade the neutralizing antibody response (49, 50). In addition, the VLPs was administered parenterally while the challenge virus was applied directly to mucosal surfaces in the nasal passage. Nevertheless, immunized animals demonstrated reduced weight loss and recovered from infection faster than control animals in a standard challenge model. At autopsy, substantially reduced pathology in the lungs of the immunized animals was apparent when compared with the controls with reduced eosinophilia and lower levels of virus encoded S protein in those few lesions present. Evidently, while mucosal surface antibody levels were insufficient to prevent infection *per se* the systemic immune response induced by the VLPs was sufficient to restrict virus proliferation and reduce the severity of disease overall. Hennrich and co-workers have described a VLP vaccine for SARS-CoV-2 based on a mini-spike form of S delivered by a VSV based replicon which also prevented weight loss and overt disease in a similar model (39) and Tan et al., have described a novel thermostable RBD decorated VLP based on the SpyCatcher technology (46) which induced very high levels of neutralizing antibody consistent with protection although a formal virus challenge was not reported. Kang et al. also reported SpyCatcher enables VLPs, with similar results (51). Plescia et al., reported SARS-CoV-2 VLPs produced in mammalian cells by co-expression of S, M, N and E (12) while Xu et al., reported mammalian cell expressed VLPs with only S, M and E consistent with the data reported here (38). Neither study investigated the immune response to the VLPs made. More recently VLPs, also made by co-expression of all four structural proteins in mammalian cells, have been used to investigate the basis of genome packaging and, in turn, the role of variant sequences on the efficiency of SARS-CoV-2 virus assembly, providing evidence that amino acid changes in N rather than S are significant drivers of variant emergence (52). In our study, VLPs were immunogenic in the absence of adjuvant and were produced using a technology that is already in use for both human and animal vaccines offering scalability and an established route to licensure (22). The VLPs described here have strong precedents in other examples derived from emerging enveloped viruses, including, among others, influenza (32, 53), Ebola (54), HIV (55), Nipah (56) and SARS-CoV (9). Their confirmation here as an effective vaccine against SARS-CoV-2 supports the case for larger scale production and clinical assessment.

## ACKNOWLEDGEMENT

We are grateful to Danielle Groves and Saskia Bakker (University of Warwick, UK) for their technical expertise and help in electron microscopy. We thank Giuseppe Ercoli and Jeremy Brown at UCL for providing us human convalescent sera and Jo Hall, Claire Ham, Adrian Jenkins, Rose Leahy and Vicky Rannow at NIBSC, all of whom provided excellent technical assistance and help for the challenge studies. We are also grateful to Professor Simon Priestnall and Alejandro Suarez-Bonnet at the Royal Veterinary College for pathology scoring and interpretation. The project was funded by UKRI-BBSRC grant # BB/V006584 to P Roy.

